# White matter pathways supporting individual differences in epistemic and perceptual curiosity

**DOI:** 10.1101/642165

**Authors:** Ashvanti Valji, Alisa Priemysheva, Carl J. Hodgetts, Alison G. Costigan, Greg D. Parker, Kim S. Graham, Andrew D. Lawrence, Matthias J. Gruber

**Author notes:** Corresponding Authors: Ashvanti Valji,; Matthias J. Gruber.

## Abstract

Across the lifespan, curiosity motivates us to learn, yet curiosity varies strikingly between individuals. Such individual differences have been shown for two distinct dimensions of curiosity: *epistemic curiosity* (EC), the desire to acquire conceptual knowledge, and *perceptual curiosity* (PC), the desire for sensory information. It is not known, however, whether both dimensions of curiosity depend on different brain networks and whether inter-individual differences in curiosity depend on variation in anatomical connectivity within these networks. Here, we investigated the neuroanatomical connections underpinning individual variation in trait curiosity. Fifty-one female participants underwent a two-shell diffusion MRI sequence and completed questionnaires measuring EC and PC. Using deterministic spherical deconvolution tractography we extracted microstructural metrics (fractional anisotropy (FA) and mean diffusivity (MD)) from two key white matter tracts: the fornix (implicated in novelty processing, exploration, information seeking and episodic memory) and the inferior longitudinal fasciculus (ILF) (implicated in semantic learning and memory). In line with our predictions, we found that EC – but not PC – correlated with ILF microstructure. Fornix microstructure, in contrast, correlated with both EC and PC, with posterior hippocampal fornix fibres - associated with posterior hippocampal network connectivity - linked to PC specifically. These findings suggest that differences in distinct dimensions of curiosity map systematically onto specific white matter tracts underlying well characterized brain networks. Furthermore, the results pave the way to study the anatomical substrates of inter-individual differences in dimensions of trait curiosity that motivate the learning of distinct forms of knowledge and skills.

## Introduction

Curiosity is described as the desire for new information that motivates seeking out and acquiring knowledge (Loewenstein, 1994; Litman, 2005). The momentary experience of curiosity (state curiosity) can be seen as an emotional-motivational state that facilitates exploration and knowledge acquisition (Silvia & Kashdan, 2009; Gottlieb & Oudeyer, 2018). Consistent with this idea, studies have shown that states of high curiosity enhance long-term memory (Kang et al., 2009; Gruber et al., 2014; McGillivray et al., 2015; Marvin & Shohamy, 2016; Stare et al., 2018; Galli et al., 2018). Furthermore, recent neuroimaging evidence suggests that state curiosity enhances memory via increased activation in the mesolimbic dopaminergic circuit including the hippocampus (Gruber et al., 2014; Kang et al., 2009). Notably, the positive effects of state curiosity on memory have been found to greatly vary between individuals in that individual variations observed in the midbrain and hippocampus activity predict the magnitude of memory enhancements (Gruber et al., 2014).

Over the last decades, between-person differences in curiosity as a personality trait (i.e. dispositional tendencies to experience and express curiosity) have been well characterized. Based on Berlyne’s (1954) suggestion that different types of curiosity are aroused by opportunities for new knowledge or sensory stimulation, trait curiosity has been split into two broad facets: curiosity as engagement with semantic knowledge - *epistemic curiosity* (EC); or as engagement with sensory stimuli - *perceptual curiosity* (PC). Building on Loewenstein’s (1994) model of aversive curiosity, Litman and colleagues further proposed that these two aspects of curiosity can be further separated into diversive/interest-based and specific/deprivation-based curiosity. Diversive/interest curiosity is linked to positive affect and is thought to energize and to direct exploration with the ultimate goal of stimulating one’s interest and reduce boredom. In contrast, specific/deprivation curiosity is accompanied by a negative, frustrated feeling of information deprivation and uncertainty, associated with a specific knowledge gap, that people are motivated to eliminate (Berlyne, 1966; Litman, 2005, 2008, Litman & Spielberger, 2003; Litman & Jimerson, 2004). Importantly, such inter-individual differences in curiosity have been found to predict job performance and academic achievement in the real world (Grossnickle, 2016; Kashdan & Yuen, 2007; Mussel, 2013).

The neuroanatomical substrates underpinning individual differences in trait curiosity are unknown. Studies investigating higher-order personality traits subsuming curiosity, however, provide a fruitful starting point to investigate the neuroanatomical connections underlying trait curiosity (DeYoung, 2014; Woo et al., 2014). For example, Privado et al. (2017) found an association between ‘openness to experience’ and microstructure of the inferior longitudinal fasciculus (ILF), a ventral, temporo-occipital association tract implicated in semantic learning and memory (Herbet et al., 2018; Hodgetts et al., 2015, 2017; Ripollés et al., 2017). Additionally, Cohen et al. (2009) showed that individual differences in novelty seeking were associated with microstructure of the fornix, a key pathway that connects the hippocampus - involved in novelty detection, exploration, information seeking and episodic memory (O’Keefe & Nadel, 1978; Kumaran & Maguire, 2009; Murray et al., 2017; Voss et al., 2017) - to the thalamus, ventral striatum, amygdala and prefrontal cortex (Saunders & Aggleton, 2007; Aggleton et al. 2015).

Here, we used multi-shell diffusion MRI and spherical deconvolution tractography to investigate whether individual differences in ILF and fornix microstructural metrics (i.e., fractional anisotropy (FA) and mean diffusivity (MD)) would be associated with individual differences in trait curiosity. Given the importance of ILF to semantic cognition (Jouen et al., 2015; Chen et al., 2017; Hodgetts et al., 2017; Ripollés et al., 2017; Herbet et al., 2018), we predicted an association between ILF microstructure and EC but not PC. In contrast, given that hippocampal circuitry supports novelty detection, exploratory behaviour and information seeking in many domains (O’Keefe & Nadel, 1978; Kumaran & Maguire, 2009; Otmakhova et al., 2013; Murray et al., 2017; Voss et al., 2017) we predicted an association between fornix microstructure and *both* EC and PC. Further, given evidence of a posterior (fine-grained) to anterior (gist-based) gradient of representational specialization along the long-axis of the hippocampus (Ranganath & Ritchey, 2012; Poppenk et al. 2013; Strange et al., 2014; Murray et al., 2017), we predicted that microstructure of fornical fibres associated with posterior and anterior hippocampus (Christiansen et al., 2017; Saunders & Aggleton, 2007) would be more strongly associated with PC and EC, respectively.

## Results

### Epistemic curiosity – but not perceptual curiosity – correlates with ILF microstructure

#### ILF FA

We conducted a series of permutation tests that investigated the relationships between trait curiosity scores and microstructure in *a-priori* selected anatomical tracts. For each permutation test, we corrected for multiple comparisons for the two subscales separately within the EC and PC scale. The first permutation test targeted ILF FA and EC. We found that bilaterally averaged ILF FA did not significantly correlate with either subscale of EC (deprivation EC, *r*(50) = 0.143, *p*_*corr*_ = 0.243, 95% CI [−0.105, 0.364]; interest EC, *r*(50) = 0.191, *p*_*corr*_ = 0.151, 95% CI [−0.0734, 0.440]). A further permutation test was conducted on bilaterally averaged ILF FA with the two subscales of PC, where again neither subscale significantly correlated with bilateral ILF FA (specific PC, *r*(50) = 0.109, *p*_*corr*_ = 0.329, 95% CI [−0.229, 0.427]; diversive PC, *r*(50) = 0.207; *p*_*corr*_ = 0.122, 95% CI [−0.109, 0.453]).

#### ILF MD

Targeting ILF MD, a permutation test revealed a significant negative correlation between ILF MD and interest EC (*r*(50) = −0.289, *p*_*corr*_ = 0.038, 95% CI [−0.489, 0.007], **Figure 1A**) and a significant negative correlation between ILF MD and deprivation EC (*r*(50) = −0.388, *p*_*corr*_ = 0.004, 95% CI [−0.572, −0.124], **Figure 1B**). In contrast, bilateral ILF MD was not significantly correlated with any subscale of PC (diversive PC, *r*(50) = 0.020, *p*_*corr*_ = 0.710, 95% CI [−0.260, 0.271], **Figure 1C**); specific PC, *r*(50) = −0.134, *p*_*corr*_ = 0.267, 95% CI [−0.392, 0.157], **Figure 1D**).

**Figure 1.**
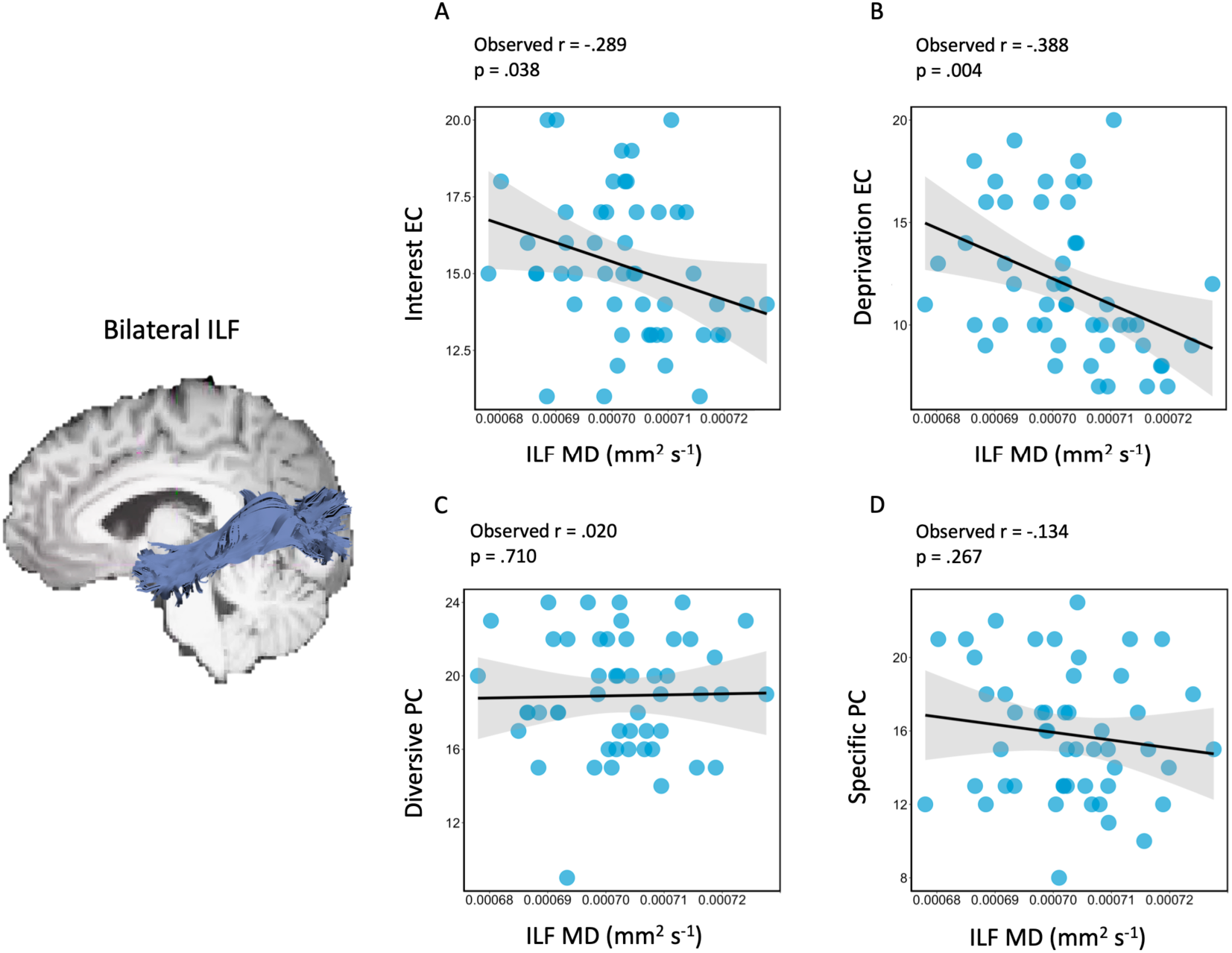
Inferior longitudinal fasciculus microstructure only shows relationship with epistemic curiosity. These results were obtained from non-parametric permutation tests that corrected for multiple comparisons across the two subscales within the Epistemic Curiosity scale (EC) and Perceptual Curiosity scale (PC). A significant negative correlation was found between MD (mm^2^ s^-1^) of the inferior longitudinal fasciculus (ILF) with interest- and deprivation EC (**A, B**, respectively). No significant correlations were found between ILF MD (mm^2^ s^-1^) with diversive and specific PC (**C, D**, respectively). The line of best fit and 95% confidence interval (CI) are shown on each scatter plot with 50 data points.

Neuropsychological and imaging evidence suggests that semantic knowledge is represented bilaterally in the anterior temporal lobes (ATL) but may show subtle inter-hemispheric (left > right) gradations for verbal stimuli (Rice et al., 2015; Hoffman & Lambon Ralph, 2018). Therefore, we asked whether the significant correlation between bilateral ILF MD and both EC subscales were driven specifically by the left as opposed to the right ILF. Separate permutation tests were conducted for each subscale of EC with left ILF MD and right ILF MD as the two variables of interest (i.e., correcting for multiple comparisons across the two hemispheres). The first permutation test on deprivation EC found that both left and right ILF MD significantly correlated with deprivation EC (left ILF: *r*(50) = −0.341, *p*_*corr*_ = 0.016, 95% CI [−0.566, −0.078]; right ILF: *r*(50) = −0.358, *p*_*corr*_ = 0.012, 95% CI [−0.564, −0.106]). The second permutation test investigating whether interest EC correlates with left and right ILF MD indicated a numerical negative relationship for both tracts but neither reached significance with the adopted multiple comparisons correction (left ILF: *r*(50) = −0.254, *p*_*corr*_ = 0.066, 95% CI [−0.491, 0.086]); right ILF: *r*(50) = −0.267, *p*_*corr*_ = 0.051, 95% CI [−0.472, −0.056]).

In order to assess whether bilateral ILF MD correlations with subsets of EC were significantly different from each other as well as subsets of PC, we conducted directional Olkin’s Z-tests (Cocor R package; Diedenhofen & Musch, 2015). For EC, we found that the correlation between ILF MD and deprivation EC was not significantly different to the correlation between ILF MD and interest EC (z(50) = 0.849, p = 0.198). Comparing EC and PC subscales, however, we found that the correlation between ILF MD and deprivation EC was significantly stronger than the correlation between ILF MD and specific PC (z(50) = 1.721, p = 0.043), and the correlation between ILF MD and diversive PC (z(50) = 2.212, p = 0.014). Furthermore, we found that the correlation between ILF MD and interest EC was significantly stronger than the correlation between ILF MD and diversive PC (z(50) = 2.407, p = 0.008), however, the correlation between ILF MD and interest EC was not significantly stronger than the correlation between ILF MD and specific PC (z(50) = 1.172, p = 0.121).

### Interest-based epistemic curiosity correlates with fornix microstructure

#### Fornix FA

Regarding fornix FA, permutation tests revealed a significant positive correlation between interest EC and fornix FA (*r*(51) = 0.281, *p*_*corr*_ = 0.039, 95% CI [−0.008, 0.491], **Figure 2A**). In contrast, deprivation EC showed no significant correlation with fornix FA (*r*(51) = 0.155, *p*_*corr*_ = 0.214, 95% CI [−0.120, 0.422], **Figure 2B**). A second permutation test was conducted on fornix FA with the two subscales of PC, diversive and specific, but neither subscale significantly correlated with fornix FA (specific PC, *r*(51) = 0.111, *p*_*corr*_ = 0.328, 95% CI [−0.266, 0.4252]; diversive PC, *r*(51) = 0.064, *p*_*corr*_ = 0.466, 95% CI [−0.204, 0.351]).

**Figure 2.**
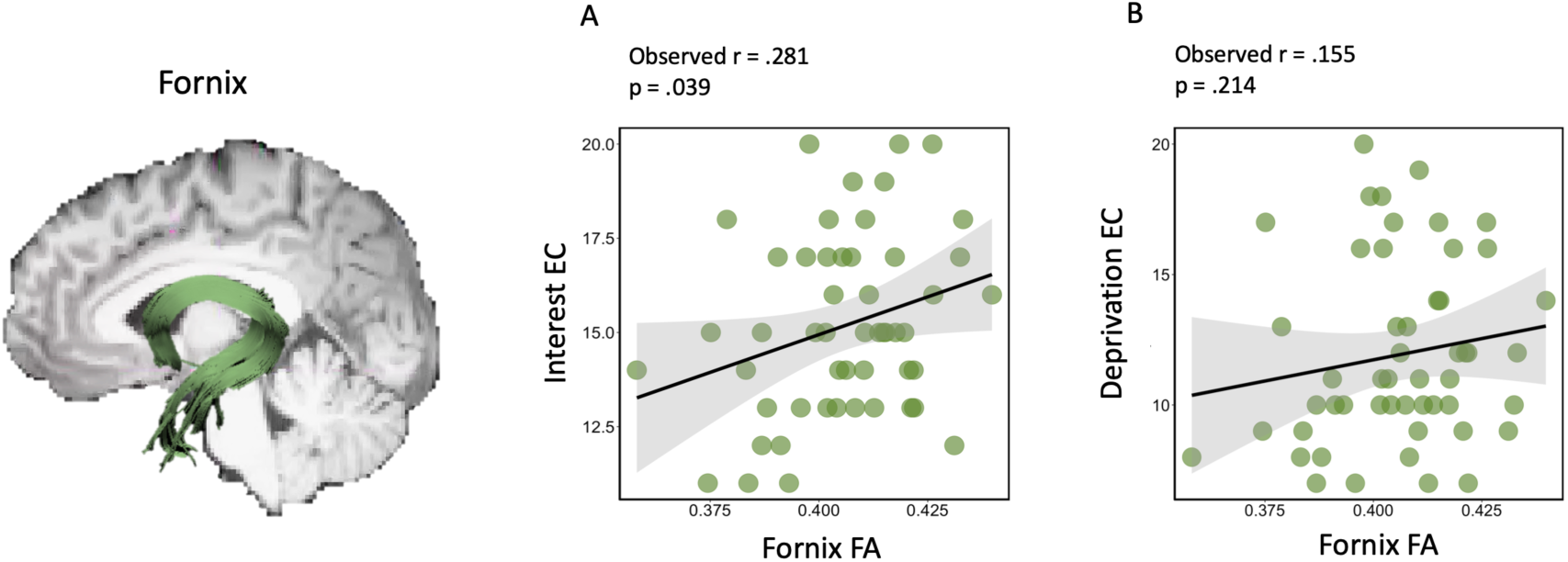
Fornix microstructure shows relationship with aspects of epistemic curiosity. These results were obtained from non-parametric permutation tests correcting for multiple comparisons across subscales within the Epistemic Curiosity scales (EC). A significant positive correlation was found between fractional anisotropy (FA) of the whole fornix and interest EC **(A)** but not with deprivation EC **(B)**. The line of best fit and 95% confidence interval (CI) are shown on each scatter plot with 51 data points.

#### Fornix MD

Despite the earlier findings of a significant positive correlation between interest EC and fornix FA, permutation tests revealed no significant negative correlation between fornix MD and interest EC (*r*(51) = −0.110, *p*_*corr*_ = 0.332, 95% CI [−0.372, 0.171]) or deprivation EC (*r*(51) = −0.029, *p*_*corr*_ = 0.574, 95% CI [−0.314, 0.296]). The second permutation test, investigating the association between fornix MD and the two subscales of PC, also showed that neither specific nor diversive PC significantly correlated with fornix MD (specific PC, *r*(51) = −0.250, *p*_*corr*_ = 0.070, 95% CI [−0.499, 0.054]; diversive PC, (*r*(51) = −0.159; *p*_*corr*_ = 0.214, 95% CI [−0.398, 0.113]).

### Specific perceptual curiosity shows an association with posterior hippocampal fornix microstructure

Recent accounts postulate a posterior-anterior gradient of representational granularity along the long axis of the hippocampus, linked to a gradient in anatomical connectivity (Aggleton, 2012; Strange et al., 2014), from ‘fine’ perceptual detail to ‘course’ or gist-like representations (Poppenk et al., 2013; Robin and Moscovitch, 2017; Sheldon et al., 2019). This account suggests that a stronger correlation might be evident between posterior hippocampal fornix and PC, and anterior hippocampal fornix and EC, respectively. To test this, we explored the relationship between *specific* PC (i.e., associated with detailed perceptual information seeking) and anterior/posterior hippocampal fornix MD. Conversely, to pinpoint how EC is associated with the anterior/posterior hippocampal fornix FA, we focussed our analyses on *interest* EC.

A first permutation test (corrected for multiple comparisons) targeted the three individual fornix segmentations (i.e., left anterior, right anterior, bilateral posterior hippocampal fornix). (Note that posterior hippocampal fornical fibres form the medial fornix cannot easily be separated into separate hemispheres). We found that specific PC significantly correlated with posterior hippocampal fornix MD (*r*(51) = −0.277, *p*_*corr*_ = 0.047, 95% CI [−0.528,0.056], **Figure 3B**), but it did not correlate significantly with left or right anterior hippocampal fornix MD (left: (*r*(51 = −0.189, *p*_*corr*_ = 0.176, 95% CI [−0.451,0.062]), **Figure 3A**; right: (*r*(51) = −0.028, *p*_*corr*_ = 0.610, 95% CI [−0.289,0.264]). This finding suggests that specific PC might mainly be supported by fornical fibres that have connections to the posterior hippocampus. Olkin’s z-tests were employed to test whether the correlation between specific PC and posterior hippocampal fornix MD was significantly different than the correlation between specific PC and left/right anterior hippocampal fornix MD. The correlation between posterior hippocampal fornix MD and specific PC was not significantly different than the correlation between *left* anterior hippocampal fornix MD (z (51) = −0.934, *p* = 0.175), however, it was significantly different than the correlation between *right* anterior hippocampal fornix MD and specific PC (z (51) = −2.268, *p* = 0.012).

**Figure 3.**
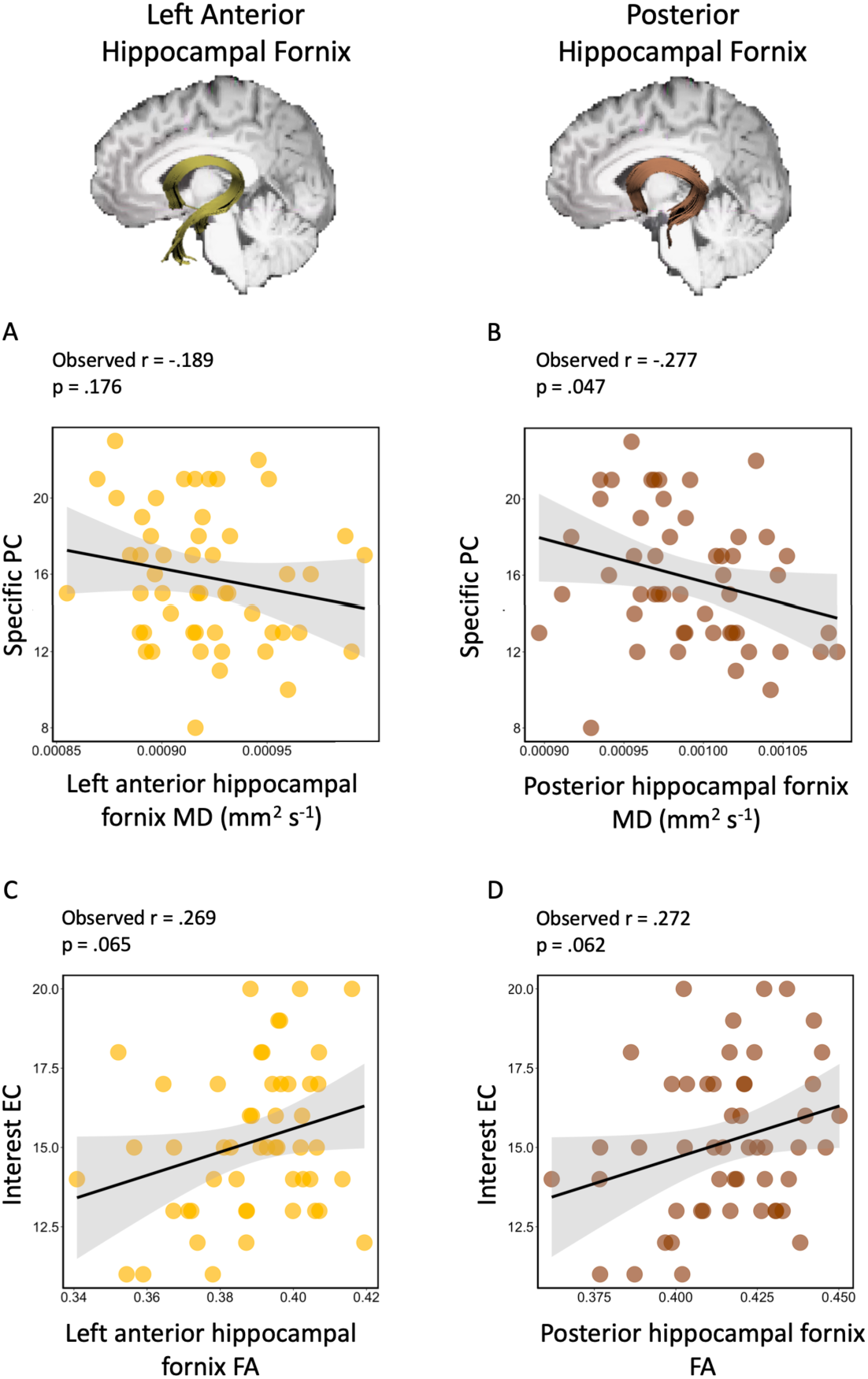
Specific perceptual curiosity, but not interest epistemic curiosity, shows a significant difference between correlations with anterior and posterior hippocampal fornix microstructure. These results were obtained from non-parametric permutation tests correcting for multiple comparisons across the three individual fornix segmentations. Specific PC did not significantly correlate with MD (mm^2^ s^-1^) of the left anterior hippocampal fornix (i.e., fornix fibres that project specifically into anterior hippocampus) **(A)** but was found to negatively correlate with MD (mm^2^ s^-1^) of the posterior hippocampal fornix **(B)**. Interest EC did not significantly correlate with FA of the left anterior hippocampal fornix (**C**), nor with FA of the posterior hippocampal fornix **(D)**. The line of best fit and 95% confidence interval (CI) are shown on each scatter plot with 51 data points.

In contrast, although we found that interest EC significantly correlates with *whole* fornix FA, the three distinct fornix segmentations did not reveal significant correlations with interest EC after correcting for multiple comparisons (left anterior hippocampal fornix FA, *r*(51) = 0.269, *p*_*corr*_ = 0.065, 95% CI [−0.029, 0.521], **Figure 3C**; right anterior hippocampal fornix FA (*r*(51) = 0.080, *p*_*corr*_ = 0.479, 95% CI [−0.161, 0.307]; posterior hippocampal fornix FA, *r*(51) = 0.272, *p*_*corr*_ = 0.062, 95% CI [−0.009, 0.479], **Figure 3D**). Olkin’s z-test indicated that the correlation between left anterior hippocampal fornix FA and interest EC was not significantly different than the correlation between posterior hippocampal fornix FA and interest EC (z (51) = 0.031, *p* = 0.488). In addition, Olkin’s z-test indicated that the correlation between right anterior hippocampal fornix FA and interest EC was not significantly stronger than the correlation between posterior hippocampal fornix FA and interest EC (z (51) = 1.443, *p* = 0.075).

In summary, we found that two individual subscales that tap into epistemic and perceptual curiosity traits showed significant correlations with fornix microstructure. In particular, we found that *whole* fornix FA significantly correlated with interest EC suggesting that interest EC relates to microstructure in fornical fibres that connect with anterior and posterior hippocampus. In contrast, specific PC specifically correlated with posterior hippocampal fornix microstructure, which was significantly stronger compared to the relationship with right anterior hippocampal fornix microstructure.

## Discussion

Curiosity motivates us to seek out information and it facilitates knowledge acquisition (Loewenstein, 1994; Litman, 2005; Silvia & Kashdan, 2009; Gottlieb & Oudeyer, 2018). While a fledgling line of research has shown that curiosity states - the momentary experience of curiosity - enhance hippocampus-dependent memory (for a review, see Gruber et al., 2019), there is also a broad spectrum of variation in *stable* tendencies to experience or express curiosity – trait curiosity. Importantly, trait curiosity has been shown to predict real-world outcomes, such as academic achievement and job performance (Kashdan & Yuen, 2007; Mussel, 2013). Here, we found that ILF microstructure correlated with both interest and deprivation EC traits, but not with PC traits. Additionally, fornix microstructure was associated with interest - but not deprivation - EC, and specific - but not diversive - PC. In particular, while microstructure of the whole fornix correlated with interest EC, specific PC correlated with posterior hippocampal fornix microstructure. These findings support the notion that curiosity is a multifaceted motivational construct and that distinct aspects of curiosity map onto specific white matter tracts underlying well-characterized brain networks that support distinct representational systems (Murray et al., 2017).

### Epistemic curiosity and ILF microstructure

The ILF, which connects ventral aspects of ATL, occipito-temporal, and occipital cortex (Herbet et al., 2018; Panesar et al., 2018), appears critical for bidirectional interactions between an ATL-based bilateral semantic ‘hub’ and representations supported by occipital and middle/posterior temporal regions (Patterson et al., 2007; Lambon Ralph et al., 2017; Chen et al., 2017). In addition to demonstrations of altered ILF microstructure in semantic dementia (Agosta et al., 2010), recent studies report associations between bilateral ILF microstructure and individual differences in semantic learning (Ripollés et al., 2017) and memory (Horowitz-Kraus et al., 2014; Hodgetts et al., 2017). Here, we found that participants with reduced diffusivity (i.e., lower MD values) in the ILF showed higher trait scores in both dimensions of EC. Critically, we found that the ILF supported both the general exploration of semantic information motivated by positive affect (EC as a feeling-of-interest) but also the search for specific information in order to close a knowledge gap (EC as an aversive feeling-of-deprivation) (Litman, 2005, 2008; Loewenstein, 1994; Lauriola et al., 2015). One explanation for this may be that perhaps the more that we learn, the more we are attuned to the gaps in our conceptual knowledge, and attending to these gaps is tension-producing and enjoyable at the same time (Loewenstein, 1994). In addition, the association between EC and ILF microstructure is in line with the literature on the higher-order personality trait ‘openness to experience’, of which curiosity is one facet (Woo et al., 2014). Privado et al. (2017) demonstrated that ILF microstructure was associated with levels of trait ‘openness’. Our findings extend this work by pinpointing that the exploration and specific search for semantic information might be one critical factor that carries the association between ‘openness’ and ILF microstructure.

### Curiosity and Fornix microstructure

The hippocampus is a medial temporal lobe structure supporting the encoding and recall of long-term memory (Burgess et al., 2002; Davachi, 2006; Eichenbaum et al., 2007; Murray et al., 2018). Given that the hippocampus has been implicated in a number of processes critical to curiosity, including exploration, reward seeking and novelty detection (O’Keefe & Nadel, 1978; Otmakhova et al., 2013; Murray et al., 2017; Kumaran & Maguire, 2009; Voss et al 2017), we investigated the relationship between curiosity and microstructure of the fornix - the principal tract linking the hippocampus with sites beyond the temporal lobe (Saunders & Aggleton 2007; Aggleton et al., 2015). Regarding the relationship between curiosity and fornix microstructure, we performed analyses targeting the microstructure of the whole fornix, but also the anterior and posterior hippocampal fornix segments that correspond to the functional subdivisions of the anterior and posterior hippocampus, respectively (Christiansen et al., 2017; Saunders and Aggleton, 2007). Given current theoretical ideas, the anterior and posterior hippocampal fornix fibres may reflect functional subdivisions of the anterior and posterior hippocampus reflecting gist-based (schematic) and perceptually detailed (episodic) information, respectively (Robin & Moscovitch, 2017; Poppenk et al., 2013; Ranganath & Ritchey, 2012; Sheldon et al., 2019). Therefore, the present study investigated whether the functional subdivisions of the fornix, connecting to the anterior and posterior hippocampus, may potentially map onto diversive/interest and specific/deprivation curiosity, respectively. Partially consistent with this hypothesis, we found that posterior hippocampal fornix (but not the anterior hippocampal fornix) microstructure (FA) showed an association with specific PC, which is described as the desire to reduce uncertainty by searching for specific novel perceptual information. Of note, recent work has highlighted a role for (posterior) HC circuitry in detailed visual exploration (Liu et al., 2017; Voss et al., 2017) and Risko et al. (2012) used a scene-viewing task to demonstrate that participants’ PC trait score predicted the degree to which they explored visual scenes. These studies using eye-movements to investigate hippocampal- and curiosity-based visual exploration and our present findings on fornix microstructure highlight how individual differences in curiosity may play a critical part in the degree of exploration of one’s perceptual environment, serving to accumulate information from the visual world, contributing to the formation of detailed memory representations mediated by posterior hippocampal circuitry.

In contrast, we found that interest EC positively correlated with microstructure of the whole fornix, rather than anterior hippocampal fornix specifically. Interest EC is described as the desire for diversive exploration and information seeking, which is accompanied by positive affect (Litman, 2008). Although interest EC reflects the reward-driven explorative search for new knowledge, presumably involving interactions between anterior hippocampal schematic or gist-based representations and reward/value representations mediated by nucleus accumbens and ventromedial prefrontal cortex (Poppenk et al., 2013; Aggleton et al., 2015), interest EC also triggers search for detailed information rather than gist-based information, presumably involving more fine-grained posterior hippocampal representations. Interest EC may therefore involve coordination along the entire hippocampal longitudinal axis, in line with the graded and overlapping nature of long axis connectivity (Aggleton, 2012; Strange et al., 2014).

### Limitations and future directions

First, our correlational analyses cannot establish causality in brain-behaviour relationships. Longitudinal studies would be needed to determine whether trait curiosity shapes white matter connections, vice versa, or whether both reinforce each other in a bidirectional manner. For instance, recent work on adaptive myelination suggests that change in myelination through activity-dependent adaptation of an initially hard-wired process occurs in response to experiences and contributes to learning (Bechler et al., 2018). Second, interpreting the biological relevance of tensor metrics from white matter tracts, such as FA and MD, can be challenging. Whilst FA and MD are typically inversely related, where a high FA and low MD suggest ‘stronger’ white matter connectivity (Vettel et al., 2017), we found that for the majority of microstructure-curiosity correlations that only one of the two diffusion metrics significantly correlated with curiosity, suggesting they are not redundant measures. Dissociations between FA and MD measures could be due to a number of biological properties such as axon diameter and density, myelination and the arrangement of fibres in a given voxel (Beaulieu, 2002). For instance, high FA has been found to reflect high myelin density and structured histological orientation whereas high values of MD are more likely to reflect low myelin density and diffuse histological orientation (Seehaus et al., 2015). Future work on the microstructural correlates of trait curiosity could apply advanced modelling techniques, such as the “Neurite Orientation Dispersion and Density Imaging” model (NODDI (Zhang et al., 2012)) for estimating biologically specific properties of the white matter.

Our study involved self-report questionnaires to measure distinct curiosity traits. While self-report questionnaires of personality have well known limitations (Vazire & Carlson, 2010), such instruments, unlike task-based measures, are designed to maximise consistent inter-individual differences, have high reliability and predict real world outcomes (Grossnickle, 2016; Eisenberg et al., 2019; Enkavi et al., 2019). Nevertheless, future studies should consider examining the link between task-evoked states of curiosity and trait curiosity, and whether the same white matter tracts mediate state effects of curiosity in different aspects of learning and memory.

### Conclusion

The present study found inter-individual variation in the microstructure of the fornix related to interest EC and inter-individual variation in the microstructure of the ILF related to both interest and deprivation EC. Furthermore, posterior hippocampal fornix microstructure was associated with specific PC. In conclusion, our findings on the relationship between curiosity traits and anatomical connections underlying well characterized brain networks provide a foundation for future studies to examine the relationship between curiosity traits, curiosity states and their neuroanatomical substrates. Our findings pave the way to further understand inter-individual differences in curiosity and which aspects of curiosity benefit language, memory and other cognitive processes cultivating a deeper knowledge and skill set.

## Materials and Methods

### Participants

Fifty-one healthy female adult undergraduate students, with a mean age of 20 years (SD ± 1, range = 19-24) participated. They provided written consent prior to participating in the study, which was approved by the Cardiff University Research Ethics Committee, and received a remuneration of approximately £25 for their participation.

### Trait curiosity measures

Participants completed the *Epistemic Curiosity Scale (EC)* (Litman, 2008) and the *Perceptual Curiosity Scale (PC)* (Collins et al., 2004), along with other measures not relevant to the current study. The EC scale consists of five interest EC items and five deprivation EC items with participants answering on a scale ranging from 1 (*almost never*) to 4 (*almost always*). The interest EC items are associated with behaviours that stimulate positive affect, or involve learning something completely new (e.g. “I enjoy learning about subjects that are unfamiliar to me”). In contrast, deprivation EC items describe behaviours that reduce negative feelings of information deprivation and uncertainty (e.g. “I can spend hours on a single problem because I just can’t rest without knowing the answer”). The PC scale (Collins et al., 2004) comprised of twelve items (6 diversive PC items and 6 specific PC items) and again participants respond on a scale that ranged from 1 (*almost never*) to 4 (*almost always*). The diversive PC items describe exploratory behaviours in which one seeks out new places and a broad range of sensory stimulation (e.g. “I like to discover new places to go”), whereas specific PC describes exploration of novel, specific and sensorially stimulating stimuli (e.g. “When I hear a strange sound, I usually try to find out what caused it”). The Cronbach’s alpha coefficients for the scales were all >= .70 suggesting good internal consistency.

### Imaging acquisition

Imaging data were obtained at CUBRIC, Cardiff University on a 3 Tesla MRI scanner (Siemens Magnetom Prisma) with a 32-channel head coil. T1-weighted structural 3D images were acquired using an MPRAGE sequence (orientation = sagittal; TR = 2250ms; TE = 3.06ms; TI = 900ms; flip angle = 9°; FOV = 256mm^2^; slice thickness = 1mm; voxel size = 1mm^3^; number of slices = 224; bandwidth = 230Hz/pixel; total acquisition time = 7 minutes 36 seconds).

Diffusion weighted images were acquired using a multi-shell sequence (orientation = transversal/axial; TR = 9400ms; TE = 67.0ms; FOV = 256mm^2^; slice thickness = 2mm; voxel size = 2mm^3^; number of slices = 80). Diffusion gradients were applied in (i) 30 isotropic directions by using a diffusion-weighted factor b=1200sec/mm^2^, (ii) in 60 isotropic directions by using a diffusion-weighted factor b=2400sec/mm^2^, and (iii) a volume without diffusion gradients (b=0sec/mm^2^) (bandwidth = 1954Hz/pixel; total acquisition time = 15 minutes 51 seconds).

### Diffusion MRI pre-processing

T1-weighted structural images were subjected to a ‘brain-tissue only’ mask using FSL’s Brain Extraction Tool (Smith, 2002). Using ExploreDTI (v4.8.3; Leemans et al., 2009) each b-value image was then co-registered to the T1 structural image. Subsequently, all b-value images were corrected for head motion and eddy currents within ExploreDTI. Tensor fitting was conducted on the b-1200 data given the tensor model assumes hindered (Gaussian) diffusion, and at lower b-values more of the signal is due to hindered rather than restricted diffusion (Jones et al., 2013). To correct for voxel-wise partial volume artefacts arising from free water contamination, the two-compartment ‘Free Water Elimination’ (FWE) procedure was applied to the current b-1200 data – this improves reconstruction of white matter tracts near the ventricles such as the fornix (Pasternak et al., 2009, 2014), yielding whole brain voxel-wise free-water corrected FA and MD tissue maps. Following FWE, corrected diffusion tensor-derived structural metrics were computed. Fractional anisotropy (FA), reflects the extent to which diffusion within biological tissue is anisotropic (constrained along a single axis). MD (10^-3^ mm^2^ s^-1^) reflects overall degree of diffusivity (Vettel et al., 2017). The resulting free water corrected FA and MD maps were inputs for the tractography analysis.

### Tractography

As higher b-values allow for better fibre orientation estimations (Vettel et al., 2017), we performed tractography on the b-2400 data using damped Richardson-Lucy spherical deconvolution (dRL-SD). Spherical deconvolution provides a direct estimate of the underlying distribution of fibre orientations in the brain and when applied to tractography leads to accurate reconstructions of the major white matter pathway, and an improved ability to describe complex white matter anatomy (Dell’Acqua & Tournier, 2018). The algorithm extracted peaks in the fibre orientation density function (fODF) at the centre of each voxel, where streamlines along the orientation of the fODF peaks were reconstructed using a step size of 0.5mm. Streamline tracts were terminated if the direction of the pathway changed through an angle greater than 45° or if the fODF threshold fell below 0.05.

In ExploreDTI, manual tractography was carried out using AND, NOT, and SEED ROI gates on colour-coded FA maps to extract specific white matter tracts. AND gates (**Figure 4** - green) were used to extract fibres that passed through the gate, NOT gates (**Figure 4** - red) were used to exclude any fibres that passed through the gate, and finally SEED gates (**Figure 4** - blue) were used as a starting point to extract fibres that passed through this gate and then to include only those fibres that then passed through any added AND gates. Manual tractography was carried out on a minimum of 15 subjects in order to calculate a tract model to perform automated tractography on all 51 data sets (Explore DTI; Parker et al., 2013). This procedure enables the construction of white matter tracts in space in which streamlines belonging to particular anatomical features of interest consistently project to distinct sub-regions, allowing the reconstruction of streamline data by observing their projected positions (Parker et al., 2013). After running the automated tractography software each tract was visually inspected, and any erroneous fibres were pruned using additional NOT gates. These tract masks from the b=2400 data were then intersected with the b=1200 free-water corrected FA and MD maps to derive free-water corrected tract-specific measures of FA and MD values for statistical analysis.

**Figure 4.**
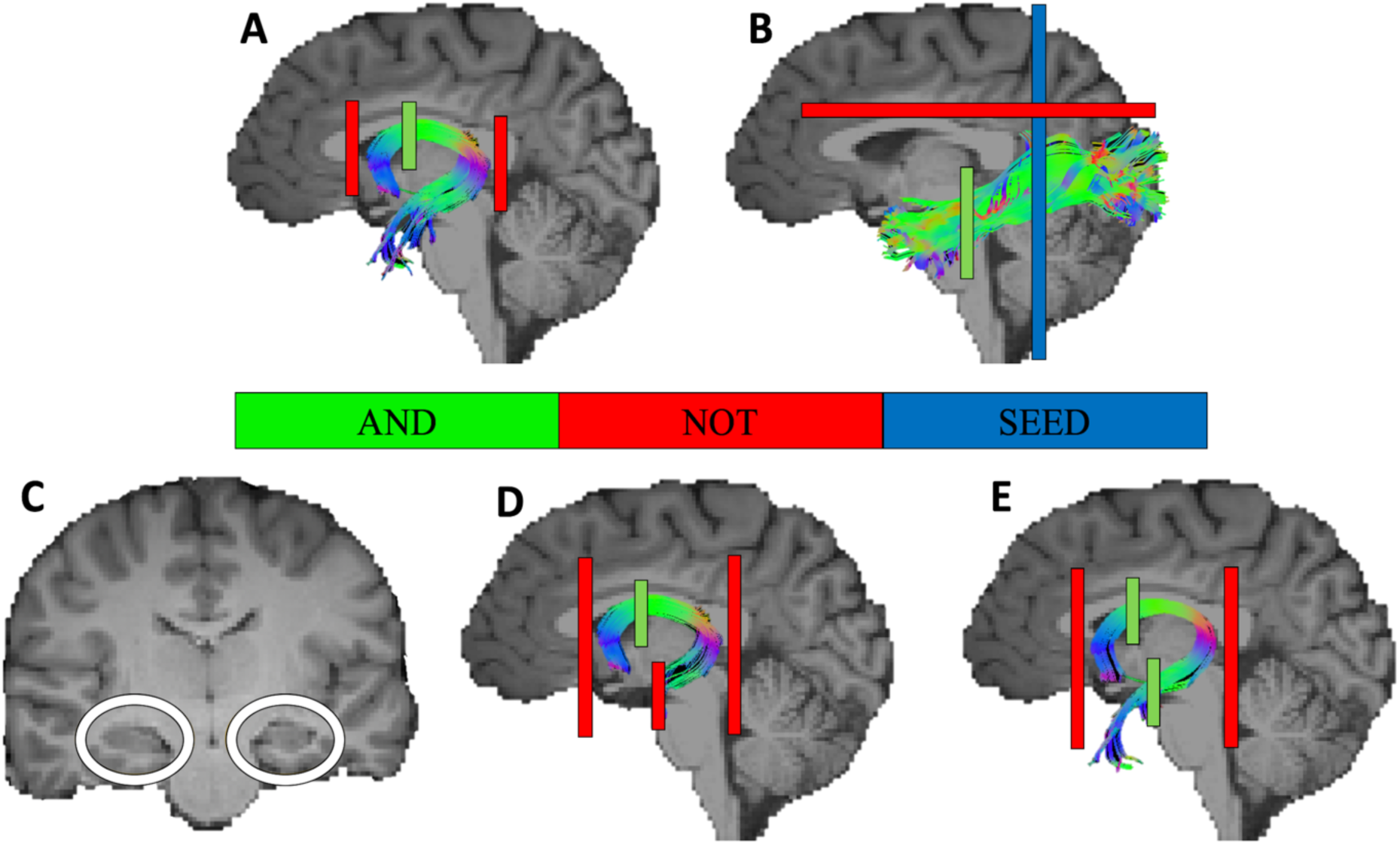
Automated tractography reconstructions of the fornix, its anterior and posterior hippocampal fornix fibres and the inferior longitudinal fasciculus (ILF). AND (green), NOT (red), and SEED (blue) ROI gates for each of the tracts are displayed on the sagittal midline plane. (**A**) Fornix tractography using AND and NOT gates. (**B**) Left ILF tractography using SEED, AND and NOT gates. (**C**) Location of AND and NOT gates for tractography of the anterior and posterior hippocampal fornix, respectively. (**D**) Posterior hippocampal fornix tractography using one additional NOT gate placed between the head and the body of the hippocampus to only include fornical fibres that connect with posterior hippocampus (i.e., hippocampal body and tail). (**E**) Anterior hippocampal fornix tractography using one additional AND gate placed between the head and body of the hippocampus (i.e., identical location as NOT gate in (D)) to include fibres that pass through this ROI gate to the anterior hippocampus.

#### Inferior Longitudinal Fasciculus tractography

The ILF (**Figure 4B**) was reconstructed using a two-ROI approach in each hemisphere (Wakana et al., 2007). In the mid-saggital slice of the brain, the coronal crosshair was placed posterior to the corpus callosum. In the coronal plane a SEED gate was drawn around the entire cortex of interest. Next in the coronal view, the last slice where the temporal lobe was separate from the frontal lobe was identified and one AND gate was drawn around the temporal lobe. Any stray fibres not consistent with the ILF pathway were removed with NOT gates. FA and MD of the right and left ILF were summed and averaged to provide a bilateral measure for the main analyses.

#### Fornix tractography

The fornix (**Figure 4A**) was traced in line with the landmarks described in Catani and Thiebaut de Schotten (2008). In the mid saggital slice of the brain, the coronal crosshair was placed at the anterior commissure and moved approximately 6 voxels posterior in the brain. In the coronal plane, one AND gate was drawn around the fornix bundle where the anterior pillars enter the body of the fornix. Finally, NOT gates were drawn around any protruding areas that were not part of the fornix.

#### Anterior and posterior hippocampal fornix tractography

In addition, we employed a method adapted from prior work to reconstruct the anterior and posterior hippocampal fornix fibres (Christiansen et al., 2017). Both anterior and posterior hippocampal fornix reconstructions required the AND and NOT gates that were applied during whole fornix tractography. Some NOT gates were augmented to enable better extraction of the anterior and posterior hippocampal streamlines of the fornix. A standard landmark for the anterior-posterior hippocampal boundary was proposed to be a small bundle of grey matter that outlines the most anterior extent of the parahippocampal gyrus that is called uncal apex or uncus (Poppenk et al., 2013). This landmark was identified for each hemisphere separately when carrying out manual tractography of the anterior and posterior hippocampal fornix. In order to perform this, the uncus was first localised at its anterior part and traced to its posterior boundary. The first coronal slice in which the uncus was not visible anymore was used as the landmark in order distinguish between fibres that project into anterior (head of the hippocampus) and posterior hippocampus (body and tail of the hippocampus) (**Figure 4C**).

After the left and right hemispheric landmarks were identified, one NOT gate on each hemisphere was drawn around the hippocampus to set boundaries for posterior hippocampal fornix tracts, removing fibres that pass through these NOT gates (**Figure 4D**). After the posterior hippocampal fornix was identified, the same coordinates of the anterior-posterior hippocampal boundary landmark for each hemisphere were used to replace the NOT gates with AND gates for the left and right anterior hippocampal fornix reconstruction (**Figure 4E**). The posterior, left, and right anterior hippocampal fornix were saved as separate tracts to aid subsequent automated tractography (**Figure 5**). Note that diffusion tensor metrics of the whole fornix and those averaged across anterior and posterior hippocampal fornix segments were highly correlated (FA, r(51) = 0.940, p < 0.001; MD, r(51) = 0.942, p < 0.001) indicating that the anterior and posterior hippocampal fornix reconstructions matched the whole fornix reconstructions.

**Figure 5.**
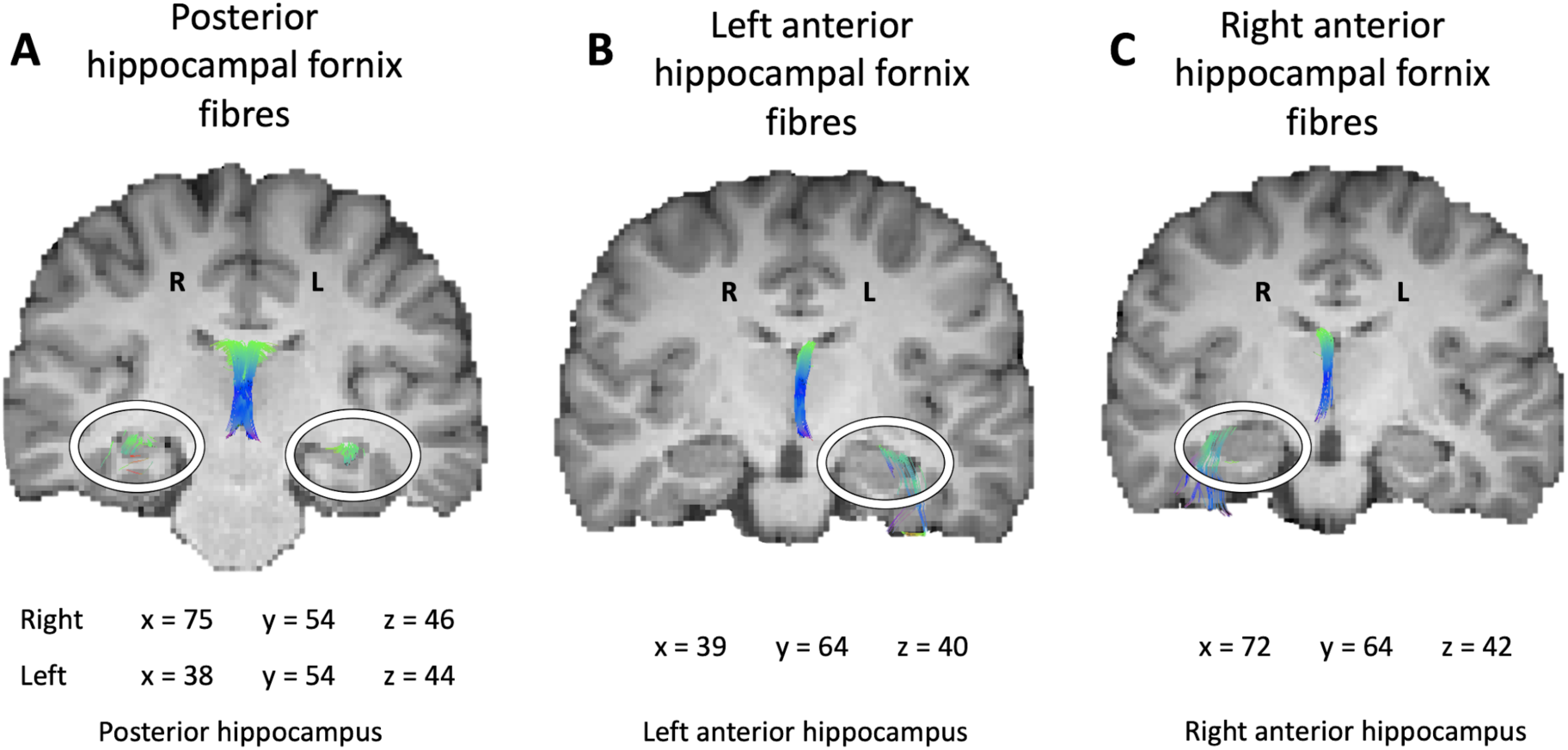
Automated tractography reconstructions of anterior and posterior hippocampal fornix fibres on coronal slices. Tractography of the fornix fibres projecting to the posterior hippocampus (**A**). Tractography of fornix fibres projecting to the left anterior hippocampus (**B**). Tractography of the fornix fibres projecting to the right anterior hippocampus (**C**).

### Statistical analyses

For the questionnaire data, in the event of missing responses (2 participants failed to give a response to one PC item), the mean value of the remaining items that were answered in the full scale was calculated which then replaced the missing item score. For each curiosity subscale (i.e., the two subscales of PC and EC), we calculated a total score for each participant. Participants’ data with diffusion tensor metrics +/– 3SD beyond the group mean were considered as outliers and removed from respective analyses. This resulted in one participant’s data being removed from all analyses involving ILF MD and a different participant’s data being removed from analyses including bilaterally averaged ILF FA.

To test for associations between curiosity trait scores and microstructure of our selected anatomical tracts, we conducted directional *Pearson’s* correlations using MATLAB. Since higher FA and lower MD is typically associated with ‘stronger’ white matter connectivity (Vettel et al., 2017), we predicted a positive correlation between levels of curiosity and FA and a negative correlation with MD.

To determine whether the *Pearson’s* correlation coefficient *r* was statistically significant, we performed non-parametric permutation tests that randomly permute the real data between participants. Permutation tests were conducted separately for the two microstructure metrics (i.e., FA and MD) and for the EC and PC scales. Importantly, we corrected for multiple comparisons across the subscales within a curiosity scale (e.g., diversive- and specific PC). The steps were as follows: First, we performed *Pearson’s* correlations on the real data (i.e., correlations between the scores of the two curiosity subscales and the microstructure measure (e.g., diversive PC with ILF MD and specific PC with ILF MD)). Thereby, we obtained the empirical correlation coefficients reflecting the relationship between the two curiosity subscales and a specific microstructure measure. Second, within each curiosity subscale, we shuffled the curiosity scores across participants, which resulted in pairs containing a curiosity score and a microstructure value that is randomly assigned across participants. On these shuffled data, we then calculated surrogate *Pearson’s* coefficients for the two curiosity subscale scores and the microstructure metric, and saved the maximum surrogate *Pearson’s r* across the two correlations (i.e., subscale-microstructure_^max^_) (Groppe, Urbach & Kutas, 2011). Third, the second step was repeated 5000 times. Based on the 5000 permutations, we created a null distribution of all surrogate subscale-microstructure_^max^_ coefficient values and determined the alpha cut-off point (*p* < 0.05; one-sided; i.e., 4750th data point of the surrogate null distribution) in order to test the statistical significance of the real *Pearson’s* coefficients reflecting the relationship between the two subscales and the microstructure measure. This approach allowed us to correct for multiple comparisons across the two subscales within each curiosity scale. In follow-up analyses for specific curiosity subscales (e.g., interest EC subscale), we also performed follow-up permutation tests that corrected for multiple comparisons across both hemispheres (e.g., left and right ILF MD). The 95% confidence intervals (CI) for each correlation was derived using a bootstrapping method based on 1000 iterations. Olkin’s z test was used for the statistical comparison of dependent correlations, as implemented in the r package ‘cocor” (Diedenhofen & Musch, 2015).

## Author contributions

A.V., K.S.G., A.D.L. and M.J.G. contributed to the conception and design of the experiment. A.V. and A.C. contributed to data acquisition. All authors contributed to data analysis and interpretation. A.V. and M.J.G. drafted the manuscript and together with C.J.H., K.S.G. and A.D.L. revised the manuscript. A.D.L. and M.J.G. jointly supervised this work.

## Acknowledgements

We would like to thank the funders of this research as well as Ofer Pasternak for providing the free-water correction pipeline, John Evans and Peter Hobden for scanning support, Samuel Ridgeway and Bethany Coad for support in data collection, Sonya Foley and Marie-Lucie Read for support in data pre-processing.

